# From deserts to forests: sequencing of the giant forest hog completes the genomic history of the African Suids

**DOI:** 10.1101/2025.11.19.689022

**Authors:** Marta Maria Ciucani, Sabhrina Gita Aninta, Nuno Filipe Gomes Martins, David A. Duchêne, Xi Wang, Ignacio A. Lazagabaster, Xiaodong Liu, Frederik Filip Stæger, Mikkel Schubert, Thomas Bøggild, Shixu He, Zilong Li, Rafael Reyna, Erik Meijaard, Thomas M. Butynski, Vincent B. Muwanika, Charles Masembe, Sylvain Dufour, Philippe Gaubert, Hans R. Siegismund, Ida Moltke, Anders Albrechtsen, Rasmus Heller

## Abstract

The giant forest hog (GFH, *Hylochoerus meinertzhageni*), the largest and most elusive wild pig, is found in a disjunct range across equatorial Africa, primarily inhabiting forests and woodlands. It is the only remaining African suid species without whole-genome data. The GFH is currently classified into three subspecies, differing in skull morphology and size. Here, we present the first whole-genome data of nine GFHs; one from Guinea and eight Uganda, representing the two geographically most distant subspecies. This genomic evidence allowed us to resolve the previously uncertain placement of the GFH among African suids as a sister group to the warthogs (*Phacochoerus* spp.). Also, we found a deep evolutionary split between the two GFH populations resulting in a level of genetic differentiation (*F*_ST_ > 0.75), which is similar to or higher than values observed among species of African wild pigs. The GFH likely diverged from warthogs around 4.6 Mya while the two study populations of GFH diverged at least 0.5 Mya and, subsequently, experienced very dissimilar demographic trajectories. The individual from Guinea had half of the heterozygosity level of the Uganda population, and lower value compared to all the African suids, probably as a result of the long term low *N_e_* and isolation. Due to the close affinity of GFH with the African rainforest biotic zone, our findings have implications for understanding the history of habitat change in Africa. They also indicate a very deep and previously underappreciated evolutionary divergence within the GFH that may have implications for its conservation.

## Introduction

The giant forest hog (GFH; *Hylochoerus meinertzhageni*, Thomas, 1904) is the largest member of the Suidae family and one of the world’s most elusive wild pigs (d’Huart 1993; d’Huart & Kingdon 2013). Despite its large size, the GFH was not described until 1904 (Thomas, 1904). The GFH is provisionally comprised of three subspecies (Kingdon 2013), differentiated on the basis of limited morphological evidence: *H. m. ivoriensis* in western Africa, *H. m. rimator* in central Africa, and the nominate subspecies *H. m. meinertzhageni* in eastern Africa. The western GFH differs from the other two subspecies in skull morphology, while the central African GFH is smaller than the eastern GFH (d’Huart 1993; d’Huart & Kingdon 2013). The geographic distribution of the GFH extends from central Kenya and Ethiopia in eastern Africa to Guinea in western Africa. It is primarily found in forests, woodlands and dense undergrowth where it plays a crucial role in these ecosystems (d’Huart & Kingdon 2013). The distribution of the GFH is probably a reflection of the range dynamics of the African forest belt, which has expanded and contracted along a latitudinal gradient during the Pleistocene, leaving relictual fragments of periodically contiguous closed canopy forest as far east as Mount Kenya in central Kenya (Thomas 1904; Plana 2004; d’Huart & Kingdon 2013).

While the GFH is classified as "Least Concern" on the International Union for Conservation of Nature’s Red List (d’Huart & Reyna 2016), hunting, habitat degradation, loss, and fragmentation due to deforestation and other human activities represent ongoing threats to this species. West Africa underwent a more than 20% loss of forest over the last decade and is predicted to experience at least 3% further loss by 2033 (Trew et al. 2024). Hunting for bushmeat is prevalent across most of the range of GFH (Jagadesh et al. 2023; Ingram et al. 2025). The elusive nature of the GFH means that our knowledge of its biology is limited (Reyna-Hurtado et al. 2023), making it challenging to predict their resilience to current ecological and anthropogenic pressures (d’Huart 1993).

Despite the growing body of genomic data for other wild pig species (Frantz et al. 2013; Lee et al. 2020; Garcia-Erill et al. 2022; Xie et al. 2022; Balboa et al. 2024), the GFH is not represented in such studies. The only genetic study to include GFH showed the phylogenetic placement of a single individual from Uganda as sister to the warthogs using a handful of genetic markers (Gongora et al. 2011). As such, the phylogeny and evolutionary history of the African suids remain contentious, with uncertain relations among fossils and extant species. There is also controversy surrounding the dating of key evolutionary events for Africa’s suids (Gongora et al. 2011; Garcia-Erill et al. 2022; Xie et al. 2022). Hence, the lack of genomic data for GFH precludes a full synthesis of the evolutionary history of African suids, as well as of a basic understanding of the evolution and phylogeography within the neglected GFH.

In this study, we generated the first whole-genome sequencing data of GFH from Uganda in eastern Africa (*H. m. meinertzhageni*) and Guinea in West Africa (*H. m. ivoriensis*). This allows assessment of the genomic diversity and population divergence between two of the three subspecies of GFH. We aimed to resolve the phylogenetic placement of the GFH within the African suids, including dating of divergence times. We also aimed to assess the genomic diversity, demographic history, and lineage divergence times for the GFH. Finally, we compared these properties across the African suids, as the GFH represents the final knowledge gap in this group, and discussed how these findings enhance our understanding of tropical Africa’s biogeography.

## Results

We generated whole genome sequencing data for nine GFH from two locations: eight from Uganda (*H. m. meinertzhageni*) and one from Guinea (*H. m. ivoriensis*). All GFH data were mapped to the domestic pig (*Sus scrofa*) reference genome and sequenced to a depth between 4x and 58x. We also included published whole genome data from other African pigs: eight red river hogs (*Potamochoerus porcus*), four bushpigs (*P. larvatus*), seven common warthogs (*Phacochoerus africanus*), three desert (Somali) warthogs (*Phacochoerus aethiopicus delamerei*), and one domestic pig (*Sus scrofa*) (Table S1). The depth of sequencing for the entire comparative dataset was between 13.9x and 101x (Figure 1a, Table S1). After mapping to the domestic pig reference and strict quality filtering of the reference genome and samples, the final dataset included 1,28 Gb sites located on the autosomal chromosomes.

**Figure 1:**
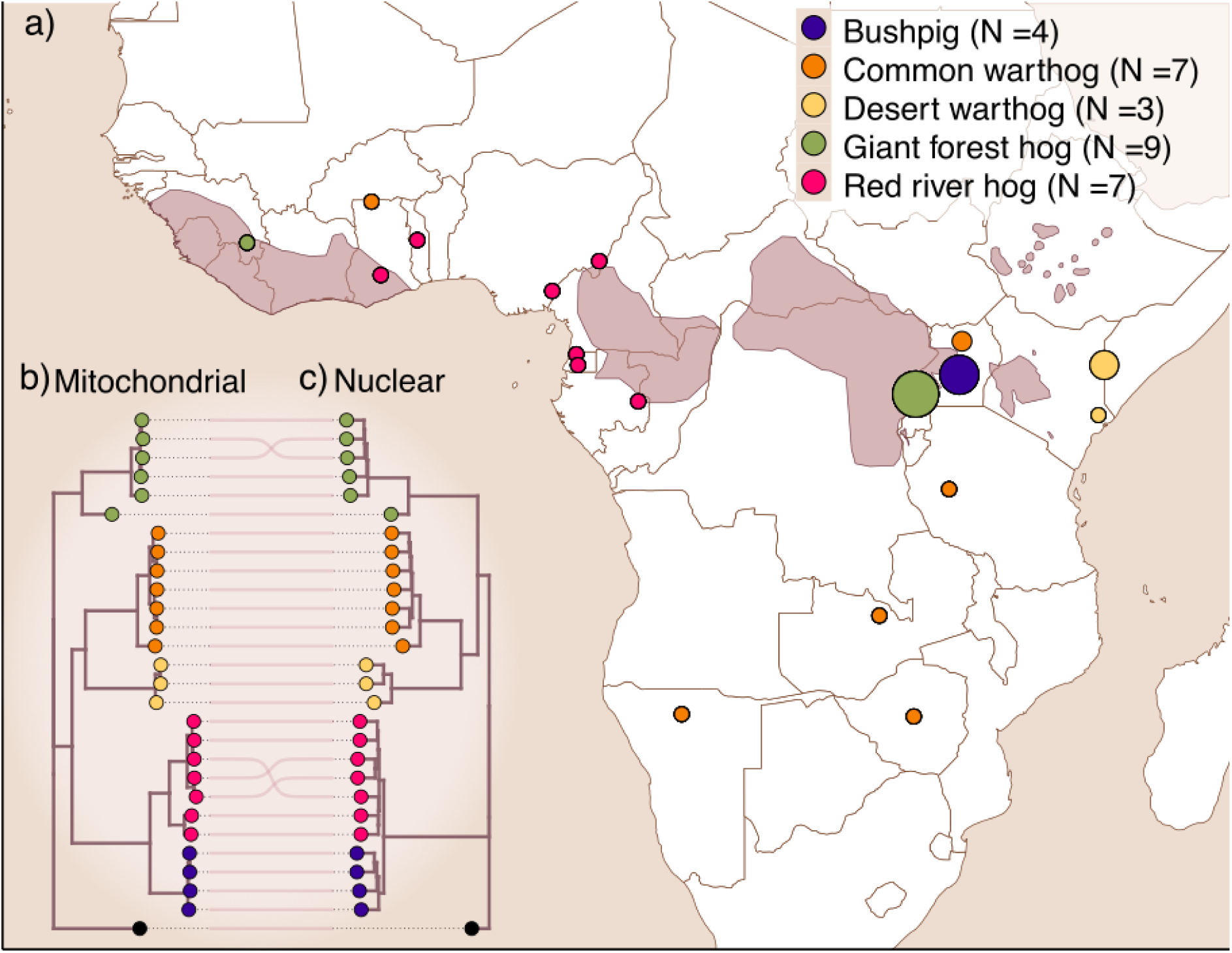
a) Sampling locations for the individuals used in this study. The geographic range of the GFH is highlighted in dark pink and was obtained from the IUCN Red List of Threatened Species (2016). b) IQtree phylogeny based on the entire mitochondrial genome (GFH individuals with depth below 5x are not included). All species split nodes have full bootstrap support. c) Species tree phylogeny generated by Astral-III estimated for 1,000 genomic regions, each 5,000 bp long. The trees in b) and c) were rooted using the domestic pig as the outgroup.

We identified closely related individuals among the GFHs from Uganda; samples 00153-00158 and 00159 and 00160 are first degree relatives. The sample with the lowest coverage (00158, 00159 and 00160) were, therefore, excluded, leaving six high coverage individuals for further analyses.

### Phylogeny of African suids

The mitochondrial DNA (mtDNA) phylogeny places the GFH basal to all other African pigs, while the GFHs from Guinea and Uganda are represented as sister taxa, with their split similar to the split between the common warthogs and desert warthogs (Figure 1b). We also explored the nuclear phylogenomic placement of the GFH samples among the diversity of the other African pigs by using Astral-III. In contrast to Figure 1b but consistent with Gongora et al. (2011), this multispecies coalescent tree, based on the autosomal data, places GFHs as sister to the warthogs (Figure 1c).

Phylogenomic dating corroborated the topology inferred using ASTRAL-III, thereby supporting the monophyly of the GFHs and the warthogs and, especially, their sister-group relationship. The dated phylogeny indicates divergence at ≈4.99 Mya (95% HPDI = 6.63-3.39) between the *Hylochoerus-Phacochoerus* (GFH, common and desert warthogs) and *Potamochoerus* (bushpig and red river hog). This is similar to the *Hylochoerus*-*Phacochoerus* divergence at ≈4.56 Mya (95% HPDI = 6.17-3.09). The common warthog and desert warthog diverged at ≈3.45 Mya (95% HPDI = 4.79-2.05), while the divergence between bushpig and red river hog is ≈1.59 Mya (95% HPDI = 2.53-0.81), similar to the one observed in the two subspecies of GFH at ≈1.75 Mya (95% HPDI = 2.77-0.81) (Figure 2).

**Figure 2:**
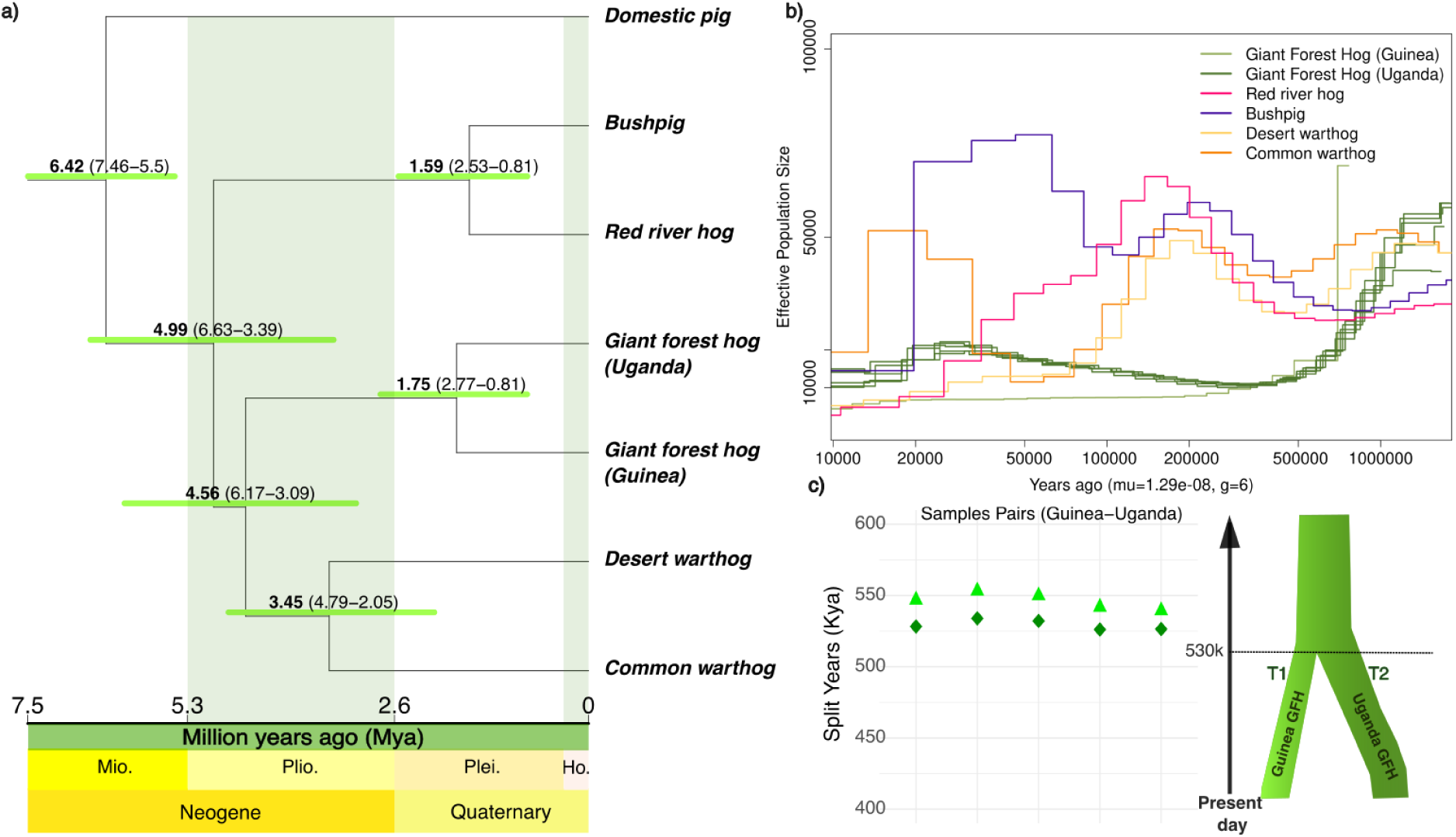
Demographic history and divergence times. a) Genome-scale dated phylogeny across species of African suids. The tree topology was inferred using IQtree and Astral. Branching times were inferred using the MCMCtree module of PAML. Green bars indicate 95% highest posterior density credible intervals (HPDI). b) Demographic history of the two GFH subspecies compared to the other African suids, inferred with PSMC. c) Two-Two (TT) method to infer the divergence time between the GFH lineage from Uganda and Guinea. A generation time of 6 years and a per-generation mutation rate of 1.29x10^-8^ were used both for PSMC and TT analyses. Triangles show the T1 split time and diamonds the T2 time.

### African suids demographic histories

To explore the demographic history of the GFH populations and place this in the context of the other African suids, we inferred the historical effective population size (*N_e_*) for the high depth samples using the Pairwise Sequential Markovian Coalescent (PSMC). GFHs from Uganda and Guinea show a sharp decrease from ca. 50k to 15k *N_e_* between 1.0 and 0.5 Mya (Figure 2b and SI). During the same time, we observed a decrease in the common warthog *N_e_*values, while the bushpig and red river hog *N_e_* were constant. After 0.5 Mya the *N_e_*trajectories of the two GFH populations separated while the other African suids *N_e_* expanded. From 120 Kya the samples from Uganda show a similar trend with their *N_e_* values increasing, peaking to ca. 20,000 around 50 Kya. Since then the *N_e_* significantly declined further around 10 Kya with values reduced to 5,000. The individual from Guinea shows a distinct trajectory in the plot with stable and low *N_e_*values (around 5,000) between 300 Kya and 50 Kya. Although we hypothesized that the demographic history of GFHs, red river hog, and bushpig might share general demographic trends, as they are all associated with forested habitats, this was not observed. *N_e_* in GFH were overall lower than for the other suid species for the majority of the last 1 million years, particularly the Guinean GFH which had a *N_e_*<10,000 individuals since 300 Kya.

Lastly, to corroborate the GFH population divergence time, we estimated the population split times between the GFHs from Uganda and Guinea using the Two-Two method (TT), which assumes a simple split with no subsequent contact. The population split is estimated to have occurred ca. 550-530 Kya for all pairs of individuals tested (Figure 2c). This result is consistent with the separation of PSMC trajectories, whereas the GFH population split times inferred with phylogenetic and population genetic methods differ considerably. Overall, however, we consistently find strong evidence that the evolutionary divergence between the western and eastern subspecies of GFH was at least 500 Kya when assuming no gene flow, and possibly considerably older if there was subsequent gene flow.

### Gene flow and genetic differentiation

To investigate potential lineage-specific gene flow among other African suids and GFH from Guinea and Uganda, we performed ABBA-BABA tests (Patterson et al. 2012). The GFH from Guinea was placed in H1, the population from Uganda was placed in H2, the outgroup was placed in H4, while all other African suid species were placed in H3 one at a time. We then tested for a difference in the amount of allele sharing between H1 and H3 compared to between H2 and H3. For gene flow detection specific to the eastern GFH we placed the common warthog and bushpig from Uganda and desert warthog from Kenya in H3. Equivalently, for gene flow specific to West African GFH, we placed a desert warthog from Kenya and red river hog from Togo in H3. No gene flow signal was found between the western and eastern GFHs and other wild African suids (Figure S1).

Subsequently, to quantify the genetic distance between the populations of GFH from Guinea and Uganda, we estimated *F_ST_*between the Guinean individual and each of the Ugandan individuals using both medium and high coverage genomes (Figure 3a). The mean *F_ST_* value between Guinean and Ugandan GFH populations was ≈0.8 (Figure 3a) indicating very high genetic differentiation and, thereby, supporting a deep split as indicated by the phylogenetic analyses. When comparing *F_ST_*among species of African suids, we found that the majority show values above 0.75 with the highest (ca. 0.9) between Guinean GFH and desert warthogs (Figure 3b). The *F_ST_*value between Guinean and Ugandan GFHs is higher than values between desert warthogs and common warthogs, and much higher than values between red river hogs and bushpigs (Figure 3b).

**Figure 3:**
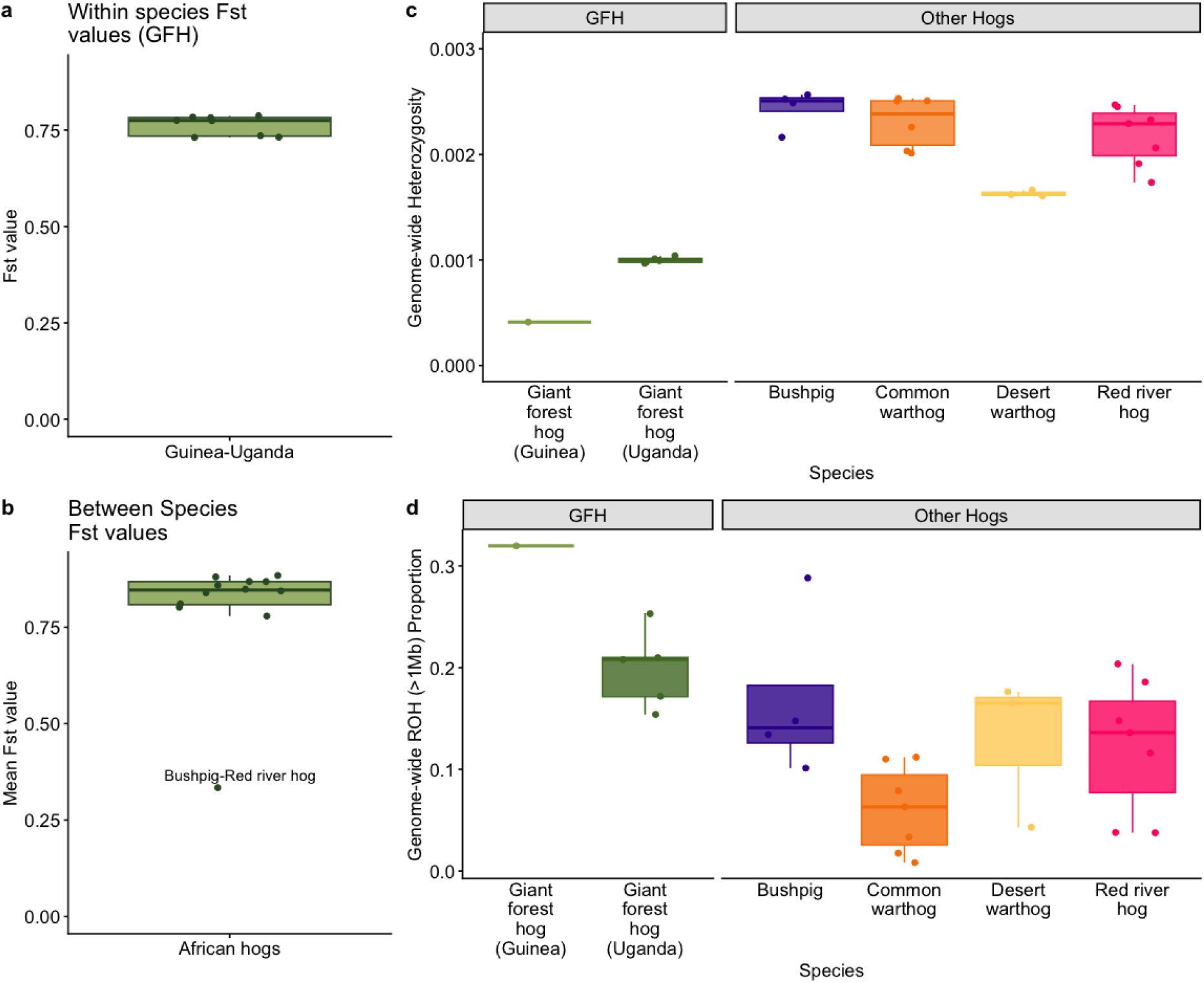
Genetic differentiation and diversity of the African suids. a) *F_ST_*between the Guinean GFH and each Ugandan GFH individual. b) *F_ST_* between all pairs of species. c) Genome-wide heterozygosity measured as the proportion of heterozygous sites per bp across each high coverage genome/individual per species. d) Proportions of genome-wide runs of homozygosity (ROHs) for all high coverage individuals by species.

We inferred the genetic diversity of GFHs by estimating the genome-wide heterozygosity. Uniform levels of heterozygosity were found among Ugandan GFHs, while the sample from Guinea displayed half as high genome-wide heterozygosity (Figure 3c). Compared with the other African suids, GFHs have much lower heterozygosity with the GFH from Guinea showing only one-third of the lowest heterozygosity estimated in other species of African suid (Figure 3c and S4a).

We then explored the runs of homozygosity (ROH) with length greater than 1Mb (ROH >1Mb) for all species. The Guinean GFH displayed the highest proportion of the genome in ROH, followed by the Ugandan GFH (Figure 3d). Overall, the common warthog shows the lowest ROH proportions among African suids, followed by the red river hog, bushpig, and desert warthog. Furthermore, the GFH from Guinea showed the highest proportions of low-medium length ROHs (<5Mb) across all individuals from all species, while keeping low proportions of long ROH (Figure S4b). This indicates that the large difference in heterozygosity between the two GFH populations is not caused by recent inbreeding in Guinea, but rather by low population sizes in recent times.

## Discussion

### Giant forest hogs show extremes in intra-specific genomic variation

In this study, we present the first genome-level analysis of the GFH from two geographical regions of the species range, thereby placing its evolutionary history within the broader context of African suid diversification. The level of genetic differentiation observed between the two GFH populations investigated here is comparable to—and in some cases exceeds—that observed between species of African suids. Remarkably, it is also greater than the genetic differentiation reported between polar bears and brown bears (Lan et al. 2022), between the two African elephant species (Pečnerová et al., in review), and among giraffe lineages (Bertola et al. 2024) now considered species (IUCN SSC Giraffe and Okapi Specialist Group Taxonomic Task Force 2025). This extremely high level of differentiation likely reflects a combination of long-standing population divergence and sustained low effective population sizes, especially in the Guinean GFH. Together, these findings suggest high amounts of genetic drift between the two GFH populations, leading to extremely high genetic differentiation on par with most of the species-level differentiation between the African suid species. Additionally, the continuously low effective population size of the Guinean GFH has led to one of the lowest genetic diversities among African ungulates, comparable to the Critically Endangered hirola (Garcia-Erill et al. 2022; Balboa et al. 2024; Bertola et al. 2024; Liu et al. 2024; Garcia-Erill et al. 2025). Notably, this low genetic diversity is primarily caused by a low effective population size over hundreds of thousands of years, and not by recent events. Therefore, ongoing human-induced habitat degradation, loss, and fragmentation, which is occurring at a higher rate in West Africa than anywhere else in tropical Africa (Aleman et al. 2018), is likely to further exacerbate an already naturally reduced genetic diversity in this region, raising concerns about genetic erosion in west African GFH (*H. m. ivoriensis*).

### Comparing African suid fossil and genetic evidence

The fossil record and evolutionary history of African suids are particularly complex. Fossils are unevenly distributed (Harris & White 1979), the material is fragmentary, and high levels of morphological variability and possible morphological convergence have complicated taxonomic assignment (Cooke 1997; Cooke, 2007; Haile-Selassie & Simpson 2013; Souron et al. 2015; Pickford and Chaïd-Saoudi, 2024). Furthermore, molecular and genomic phylogenies sometimes indicate divergences among lineages that predate the earliest African fossils (Gongora et al. 2011), suggesting mismatches between paleontological and genetic records or other inconsistencies (Garcia-Erill et al. 2022). Our phylogenomic analysis corroborates previous findings based on limited genetic markers that placed the Hylochoerini and Phacocoerini tribes as a monophyletic sister group to the Potamochoerini tribe (Gongora et al. 2011). Interestingly, the diversification between Potamochoerini and Hylochoerini-Phacochoerini, and the subsequent one between Hylochoerini and Phacochoerini appear to have occurred in rapid succession, both occurring ≈4.5-4.9 Mya and with almost identical confidence intervals (6.63-3.39 Mya and 6.17-3.09 Mya, respectively). It remains unclear, however, whether these presently African lineages diverged outside or inside Africa. Fossil evidence suggests that the major African pig lineages may have originated in other continents and arrived in Africa at different times (Pickford 2006; Arribas and Garrido 2008; Pickford 2012; Kumar and Gaur 2013). The genomic analyses presented here indicate a surprisingly deep evolutionary diversification between the two GFH populations comparable to that between the bushpig and the red river hog. Our phylogenetic and population genetic divergence time estimates of ≈1.75 Mya (2.77-0.81 Mya) and ≈0.5 Mya, respectively, with the discrepancy possibly due to either post-divergence gene flow or the difference between population divergence and genomic divergence, as discussed for warthogs in Garcia-Erill et al. (2022). Unfortunately, this deep divergence of two GFH lineages cannot be verified by fossil evidence due to the limited representation of fossil GFH in African records (Assefa et al. 2008; Lazagabaster et al. 2018; Lazagabaster et al. 2021; Rosas et al. 2022).

### Biogeographical and Afro-tropical climatic context

Among the African plant and animal diversity, many species show a west-east divergence, where populations or subspecies exhibit genetic and morphological differences along this geographic gradient (Hewitt 2004; Lorenzen et al. 2012; Couvreur et al. 2021).

Fluctuations between arid and humid climate periods during the Plio-Pleistocene and, consequently, changes in tree coverage, are proposed as the main drivers behind the west–east differentiation leading to species-level divergence (Mouline et al. 2008; Allen et al. 2021). These cycles of forest expansions and contractions, linked to glacial and interglacial phases, led to the formation of forest refugia during the Pleistocene glacial periods. The reduced distribution of closed forest in Africa during dry periods and the expansion of deserts and savannas likely represented physical barriers to forest-adapted species, leading to their differentiation into isolated populations, subspecies, and occasionally species (Cowling et al. 2008; Dupont 2011; Lorenzen et al. 2012; Allen et al. 2021).

Our inferred demographic history suggests that the GFH had large effective population sizes, comparable to the warthogs, until around 1 Mya, after which the population became severely reduced over the next 500 Ky while the other African suids increased their effective population sizes. This period, the mid-Pleistocene, marks the transition from previous 41 Ky low-amplitude cycles of climatic fluctuations to a new regime of 100 Ky higher-amplitude cycles of climatic fluctuations in tropical Africa—along with a trend towards drier climates (Dupont et al. 2001; deMenocal 2004). The ensuing continental-scale decline in closed-canopy forest would have favored more generalist suid species (bushpigs and red river hogs), and those specialized to open habitats (warthogs), and is consistent with a general trend in the fossil record of a decline in forest-adapted taxa (deMenocal 2004; Lauer et al. 2023). In GFH, our demographic analyses suggest that the long-standing demographic decline coincided with the general drying of the African climate culminating in the divergence of the two GFH ancestral populations. This might have coincided with Marine Isotope Stages (MIS) 16 and 12 (approximately 676–621 Kya and 474–427 Kya, respectively), two of the most extreme glacial periods in the mid-Pleistocene which likely accelerated the transition to drier conditions and the expansion of savannas in tropical Africa (Lisiecki and Raymo 2005; Jouzel et al. 2007; Lang and Wolff 2011; Yu and Chen 2011). Interestingly, GFHs carry a number of recently evolved cranial adaptations that suggest a recent and incomplete adaptation to a diet of low-level herbage and grasses, consistent with a scenario in which the fragmentation of previously more contiguous forests presented an evolutionary challenge to the GFH leading it to evolve adaptations to ecotones (d’Huart & Kingdon 2013). These and the following glacial periods (MIS10 and MIS8) caused major vicariance events for many African species, such as African buffalo, baboon, giraffe, warthog, lion, and spotted hyena (Rohland et al. 2005; Zinner et al. 2009; Smitz et al. 2013; Bertola et al. 2016; Coimbra et al. 2021; Garcia-Erill et al. 2022). In support of this scenario, many arid- and forest-adapted species seem to follow the pattern of expansion and contraction of forest refugia in tropical Africa, including rodents (Mouline et al. 2008; Bohoussou et al. 2015), bats, frogs, birds (De Klerk et al. 2002; Klerk et al. 2002; Herkt et al. 2016; Penner et al. 2019), and hyrax (Oates et al. 2022). Even chimpanzees (*Pan troglodyte*), seem to have followed the range contraction and expansion of forest (Hey 2010; Barratt et al. 2021; Ernst et al. 2025) making the main refugia in West Africa a reservoir of high levels of diversity and endemism (Leaché et al. 2019; Leaché et al. 2020; Ernst et al. 2025).

### Conclusions and management recommendations

We provide the first whole genome sequencing data from the GFH. Our findings show that the western African GFH in Guinea and eastern African GFH in Uganda (*H. m. ivoriensis* and *H. m. meinertzhageni,* respectively) form distinct Evolutionary Significant Units. These findings represent an initial step toward understanding the diversity within the GFH complex, as we had a small number of samples and lacked the currently recognised subspecies in Central Africa (*H. m. rimator*) and in the eastern extreme of the species’ range in the Kenya highlands (*H. m. meinertzhageni*). Nevertheless, this study underscores the need to clarify the genetic structure and evolutionary lineages of the entire *Hylochoerus meinertzhageni* complex. Although the GFH is currently classified as “Least Concern” on the IUCN Red List, the notably low level of genetic diversity observed in the Guinea population is expected to be further exacerbated by ongoing anthropogenic threats such as hunting and habitat degradation, loss, and fragmentation. More genomic research on this population will help with a reassessment of the population in Guinea. This is a research priority as the conservation status of distinct ESUs within the GFH may be more precarious than currently assumed.

## Methods

### Sample collection

The GFH sample from Guinea was collected in 2011 at the Mamou bushmeat market, located in the southwestern region of the country, during a survey conducted by SYLVATROP. Sampling followed a strictly opportunistic approach, depending on the species available at the market on a given day. The tissue sample was stored at 4°C in 90%EtOH until lab processing.

Samples from Uganda were collected as biopsies from free ranging animals at different localities within Queen Elizabeth National Park. Four samples were obtained in 2000, three in 1999 and one in 2004. All samples were stored in 25% dimethylsulfoxide (DMSO) in saturated sodium chloride (Amos and Hoelzel 1991) at ambient temperature in the field and at -80°C in the laboratory until processing.

### DNA extraction, and sequencing

Tissue samples were extracted using QIAGEN DNeasy Blood and Tissue Kit (QIAGEN, Valencia, CA, USA) following the manufacturer’s protocol. Subsequently, RNase was added to the extracted DNA to ensure the presence of only RNA-free genomic DNA. The concentration of the DNA was measured using Qubit 2.0 Fluorometer and a Nanodrop and gel electrophoresis was subsequently used to determine the quality of the genomic DNA. The samples were then sequenced using Illumina paired-end 150bp reads. Three samples from Uganda were sequenced at low depth (≈4x depth of coverage) and the remaining six samples from Uganda and Guinea were sequenced at medium-high depth (15-58x).

### Dataset

Publicly available data from the 7 common warthogs, 3 desert warthogs, 7 red river hogs, 4 bushpigs and one domestic pig were also downloaded and used in this study (Table S1) (Garcia-Erill et al. 2022; Balboa et al. 2024).

### Mapping

Sequence data pre-processing and mapping was carried out using the PALEOMIX pipeline, rev. 36ffe7c (Schubert et al. 2014). Briefly, we trimmed reads for adapter sequences using AdapterRemoval v2.3.3 (Schubert et al. 2016), merging reads overlapping at least 11 bp with the “--collapse-conservatively’ option, that masks as ’N’ mismatching, overlapping bases of the same quality. Quality trimming was not performed at this stage. Processed reads were mapped using the BWA 0.7.17 ’mem’ algorithm (Li 2013) on the Pig (*Sus scrofa*; Sscrofa 11.1; GCF_000003025.6) and Common warthog (*Phacochoerus africanus*; GCA_016906955.1) genomes. The resulting alignments were processed using SAMtools v1.11 (Danecek et al. 2021) commands “*calmd*”, “*fixmate*”, and ‘*sort*”, to update mismatch and mate meta-data, and to sort the alignments. Duplicate sequences were flagged using PALEOMIX command “*rmdup_collapsed*” for merged reads, and SAMtools “*markdup*” for all other reads. The results were merged to produce unfiltered BAM files.

Following this processing by PALEOMIX, we filtered unmapped reads, reads with unmapped mates, reads flagged as PCR duplicates or QC fail, and secondary and alternative alignments. We furthermore filtered reads with estimated insert-sizes outside the range 50 to 1000 bp, with fewer than 50 bp (or fewer than 50%) of query bases aligned to reference bases, i.e. excluding indels. Finally, we filtered pairs of reads mapping to different contigs or in the wrong orientation, and the mates of reads filtered by any of the above.

### Reference genome and site quality filtering

After mapping, we performed a variety of quality control analyses to ensure the quality of our data and reduce the impact of repetitive regions and depth of coverage on the downstream analyses for the giant forest hog populations. The *Sus scrofa* genome has RepeatMasker annotation and repeat regions where masked to avoid mismapping. Except for the mitochondrial analysis, non-autosomal chromosomes were also excluded from downstream nuclear analysis. We then estimated the global depth (read count across samples) per site separately between the low and high coverage samples using ANGSD (Korneliussen et al. 2014) applying the following parameters: “*-doCounts 1 -doDepth 1 -dumpCounts 1-maxdepth 5000 -minQ 30 -minMapQ 30*”.

The global depth distributions were then visually inspected, and we then calculated that the upper and lower 5% percentile of sites would be excluded. The sites between 5% and 95% percentile threshold were used for downstream analysis. A final sites file (.bed) was generated using bedtools (Quinlan and Hall 2010) intercept to select the overlapping sites between the filtered sites from the global depth and the repetitive regions filters.

### Mitochondrial Phylogeny

To confirm the taxonomic placement of the giant forest hog samples in our study, we performed a phylogenetic analysis on the mitochondrial genome. After mapping to the *Sus scrofa* reference genome we generated mtDNA consensus sequence for each high coverage sample using ANGSD, by including the consensus base (“*-doFasta 2*”) at each position. After calling consensus sequences, all the newly generated FASTA sequences were aligned using MAFFT v7.505 (Katoh and Standley 2013). Trimal v1.4 (Capella-Gutiérrez et al. 2009) with the parameter “*-gappyout*” was used to remove gaps and the resulting alignment was used as input in IQtree v2.1.4 (Minh et al. 2020), to perform the phylogenetic analysis. We used 1000 bootstrap replicates (“-*B 1000*”) with UFBoot2 (Hoang et al. 2018).

The option “*-m TEST*” was used to integrate ModelFinder (Kalyaanamoorthy et al. 2017) in the analysis, search for the best substitution model and perform the remaining analysis using the model. According to the BIC criteria the best fit model for our sequences was TIM2+F+I+G4

### Nuclear Phylogeny

For each sample, a majority-count consensus sequence was generated in ANGSD v0.940 using the “*-doFasta 2*” option. Bases with quality lower than 30 and reads with mapping quality lower than 30 were discarded (“*-minQ 30 -minmapq 30*”). The minimum depth for each site for each individual was set to 3x (“-*setminDepthInd* 3”), and the following additional filters were used: “*-doCounts 1 -remove_bads 1 -uniqueOnly 1 -baq 2 -C 50*”. From these consensus sequences, 1000 random regions of 5k bp long were used to generate 1000 independent gene trees using IQtree v.2.1.2. For this analysis, 1000 bootstrap replicates (“-*B 1000*”) with UFBoot2 (Hoang et al. 2018) 1000 bootstrap replicates for SH-aLRT (“*-alrt 1000*”) and *ModelFinder Plus* (Kalyaanamoorthy et al. 2017) to identify the best evolutionary model for each region. Afterwards, these region trees were concatenated to generate a species tree using ASTRAL-III (Zhang et al. 2018) with default parameters, and visualised using FigTree v1.4.4 (http://tree.bio.ed.ac.uk/software/figtree/).

### Genotype calling

We used BCFtools v1.13 (Danecek et al. 2021) to call genotypes for the African wild pig samples of high depth, including 6 giant forest hogs, 4 desert warthogs, 7 common warthogs, 4 bushpigs and 7 red river hogs. We calculated genotype likelihoods using BCFtools mpileup with the “*--per-sample-mF*” flag and performed single sample calling using BCFtools call with the “*--multiallelic-caller*” and “-G -” flags. Genotype calling was based on reads with a minimum mapping quality of 30 and bases with a minimum quality score of 30. Multiallelic sites and indels were filtered out in the called variants. We also performed genotype calling for one domestic pig, which served as the outgroup, with the same settings and merged the resulting VCF file with the one including the African wild pigs.

### D-statistics

We only kept SNPs that are polymorphic in the African wild pigs and are inside the genomic regions that passed site quality filtering in the final VCF file. We further masked genotype calls with depths below 10X and heterozygous genotypes supported by fewer than two reads in either allele. Based on this filtered genotype dataset, Admixtools2 (Maier et al. 2023) was used to calculate the D-statistics for medium-high depth individuals (> 10x). The domestic pig was used as an outgroup and the D-statistics were estimated for two scenarios: a) gene flow between the GFH from Guinea and suids from West Africa; b) gene flow between GFH from Uganda and suids from East Africa.

### Demographic history inference

#### Molecular dating

Phylogenomic dating was performed in a Bayesian framework using evenly-spaced 100 Kb windows taken from each Mb across species of African suids. Given widespread uncertainty in fossil placement in the phylogeny, we included a single calibration on the root corresponding with evidence of the earliest suine of genus *Sus* (*Sus arvernensis)* (van der Made Jan 1989) and of African hogs (*Kolpochoerus deheinzelini*) (Brunet and White 2001). Accordingly, we calibrated the root age using a uniform distribution bounded between 5.5 and 7.5 My, with a hard minimum bound and a soft maximum bound with a probability of 0.05. Molecular dating was performed on the species tree inferred as above, first calculating the gradient and Hessian matrix at the maximum likelihood estimate of molecular branch lengths (dos Reis and Yang 2011). We then ran MCMC sampling in a chain with 1,000,000,000 steps sampling every 100,000 and after a 10,000,000 step burn-in, implemented in the MCMCtree module of PAML v4.9 (Yang 2007). Model priors included gamma-distributed rates across branches and a birth-death branching process, with a GTR+Γ model for DNA substitution. We processed and visualized the output using the R packages ape (Paradis et al. 2004; Paradis and Schliep 2019) and MCMCtreeR (Puttick 2019).

#### Mutation rate

Previous demographic studies in pigs and wild suids have employed a range of mutation rate estimates. A commonly used rate of 2.5 × 10⁻⁸ mutations per site per generation, based on the human mutation rate, was applied in several genomic analyses of suids (Groenen et al. 2012; Xie et al. 2022). Other studies adopted a phylogenetically inferred rate of 2.48 × 10⁻⁹ mutations per site per year, derived from comparative analyses across wild ruminants (Chen et al. 2019; Garcia-Erill et al. 2022; Balboa et al. 2024). More recently, a pedigree-based estimate specific to domestic pigs was reported by Bergeron and colleagues (Bergeron et al. 2023), yielding a modeled mutation rate of 1.05 × 10⁻⁹ mutations per site per year. Assuming a generation time of 6 years for wild pigs (Pacifici et al. 2013), this corresponds to a per-generation rate of 0.63 × 10⁻⁸, which is approximately 2.4 times lower than the rate previously used in most suid demographic studies.

In this study we used the posterior median from the dated phylogenetic tree to estimate a new mutation rate for the African wild suids. We estimated a rate of 2.14 × 10⁻⁹ mutations per site per year which corresponds to 1.29 × 10⁻⁸ mutation rate per generation.

#### PSMC and heterozygosity

We estimated population size history for the six high coverage forest hog samples using the Pairwise Sequentially Markovian Coalescent model implemented in PSMC v0.6.5 (Li and Durbin 2011). As recommended, we generated a diploid sequence per sample using BCFtools v1.21 (Li et al. 2009), calling genotypes with -c on sites with allele support > 1, mapping quality ≥ 25, and base quality ≥ 30. Moreover, the sites needed to pass the quality filters described above and a sample-specific depth filter was calculated as depth > mean depth / 2, but minimum 10, and depth < mean depth × 1.5 (Table S2).

PSMC was run with the parameter pattern -p "4+25*2+4+6", maximum 25 iterations, and initial theta/rho ratio of 5. For scaling we used a generation time of 6 years and mutation rate of µ = 1.29×10−8 per generation. Heterozygosity was calculated as the number of heterozygous sites / total number of sites using the same filters as for psmc described above.

#### Split time estimate with the TT method

Population split times between the GFH from Uganda and Guinea were inferred using the Two-Two (TT) method, an approach estimating separate split times from a common ancestor for two populations (Schlebusch et al. 2017; Sjödin et al. 2021). For this analysis, we used the unfolded 2d-SFS from high depth individuals (> 10x coverage), polarized against the common warthog, desert warthog, red river hog and bushpig. We only used sites where all outgroups were fixed for the same allele. The same mutation rate and generation time were used to scale split times as in PSMC.

#### Population differentiation: *F_ST_* estimation

We calculated the individual-level site allele frequency (SAF) in ANGSD using GATK genotype likelihood model (*-GL 2*) with minimum mapping and base quality (*-minQ 30* and *-minMapQ 30*). From this we calculated the individual pairwise two dimensionals site frequency spectrum (SFS) using winSFS v0.7.0 (Rasmussen et al. 2022) and the genome-wide *F_ST_* using Hudson’s estimator (Bhatia et al. 2013).

We also estimated per population *F_ST_* value using called genotypes on the high depth samples for all species in this study to compare the level of differentiation across species.

For this task we used the called genotypes as input in Plink2 v2.00 (Chang et al. 2015) with the option “*--fst CATPHENO method=hudson*” and *--*“*within*” to specify the population level of each individual in a tab-separated file.

#### Runs of homozygosity (ROH) analysis

We inferred runs of homozygosity (ROH) on the high coverage samples for each species in the dataset. We filtered the samples for SNPs with a minimum MAF of 0.05 and allowed for 5% missingness. To detect runs of homozygosity, we used Plink v1.9 (Purcell et al. 2007) using the option “*homozyg*” on the autosomal chromosomes. We applied default settings and a maximum of 3 heterozygous SNPs and 20 missing calls within a scanning window. We inspected the ROHs by plotting them with proportion of heterozygous sites, SNP density in sliding windows and SNP calls along the autosomal chromosomes using a custom R script.

## Acknowledgments

M.S. was supported by the Novo Nordisk Foundation grant (NNF) NNF23SA0084103 for the Center for Basic Metabolic Research, University of Copenhagen. D.A.D. was supported by a Data Science - Emerging researcher award from NNF (NNF23OC0084647). I.M. and X.L. were funded by a Carlsberg Semper Ardens: accelerate grant (CF20-0539). Faysal Bibi from the Museum für Naturkunde in Berlin, Germany for input on fossil dating. M.M.C, F.F.S.,T.B., S.H, Z.L, and A.A were supported by NNF (NNF20OC0061343).

## Authors contribution

MMC: methodology, analysis, writing – original draft; SGA: analysis; NFGM: analysis; DD: methodology, analysis; XW: analysis; IAL: methodology, writing – review & editing ; XL: analysis; FFS: analysis; MS: methodology, analysis, writing – review & editing; TB: analysis; SH: analysis; ZL: analysis; RR: writing – review & editing, EM: writing – review & editing; TMB: writing – review & editing; VBM: sampling; CM: sampling; SD: sampling; PG: sampling, writing – review & editing; HRS, IM, AA, RH: supervision, writing – original draft, review & editing. All authors proofread and approved the final version of the manuscript.

