## Supplementary Figures for "From deserts to forests: sequencing of the giant forest hog completes the genomic history of the African Suids"

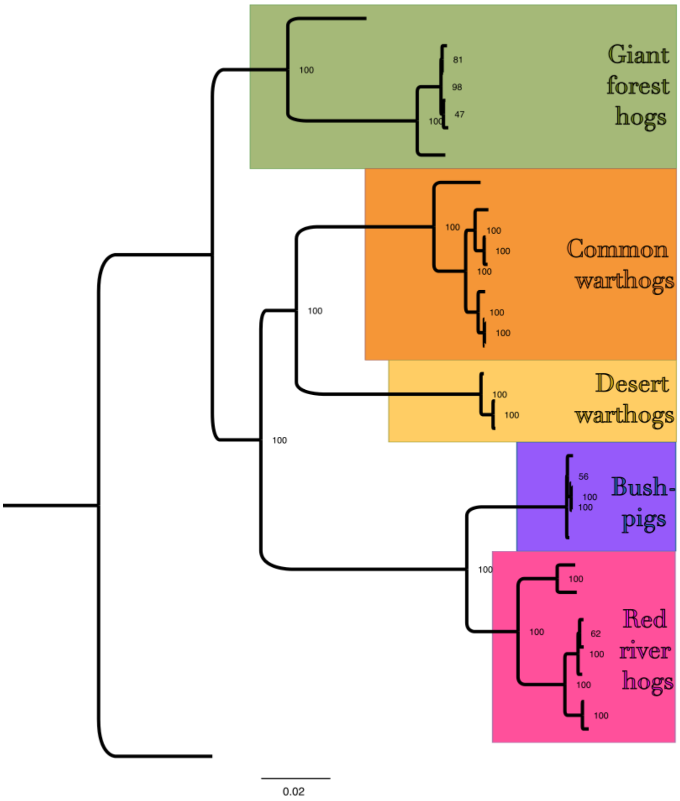


**Figure S1.** Phylogenetic tree based on the entire mitochondrial genome (GFH individuals with depth below 5x are not included) estimated using IQtree. Species split nodes have full bootstrap support (100).

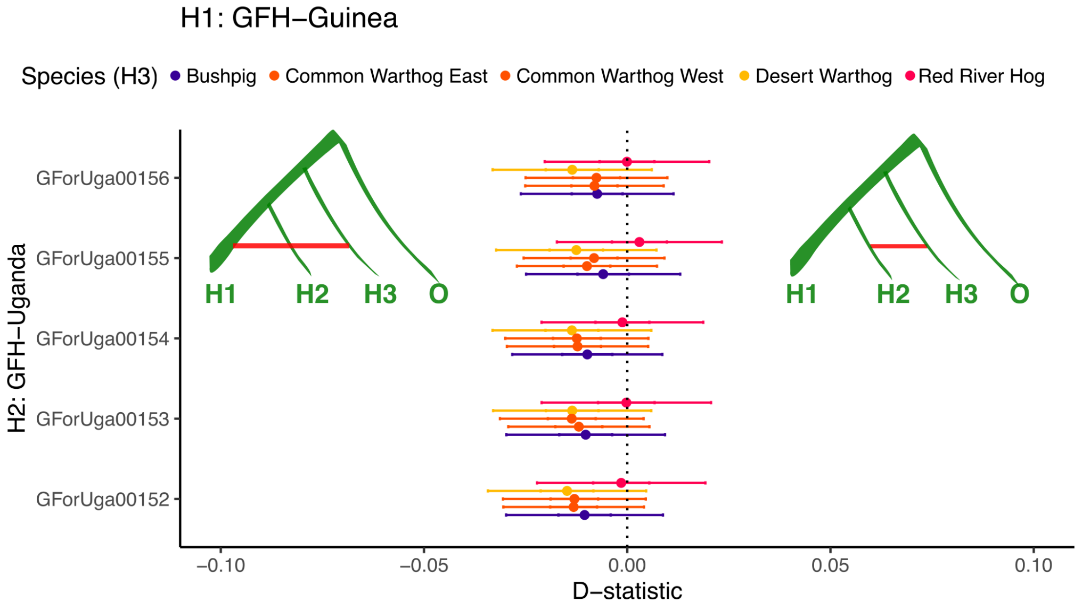


**Figure S2.** D-statistics analysis using medium-high depth (10×) samples with the domestic pig as an outgroup (O), constructed as D(H1:GFH-Guinea, H2:GFH-Uganda, H3, Domestic pig). Data are presented as the estimated D-statistic (dot) ± standard errors (error bars). On the left side of the panel, significant negative values would indicate gene flow between the GFH from Guinea (H1) and H3, represented by bushpigs, red river hogs, desert warthogs and east or west common warthog populations. Conversely, on the right side of the panel, significant positive values would indicate gene flow between GFH from Uganda (H2) and the rest of the African suids. All values and error bars are non significant, indicating no gene flow between any GFH populations and other African suids.

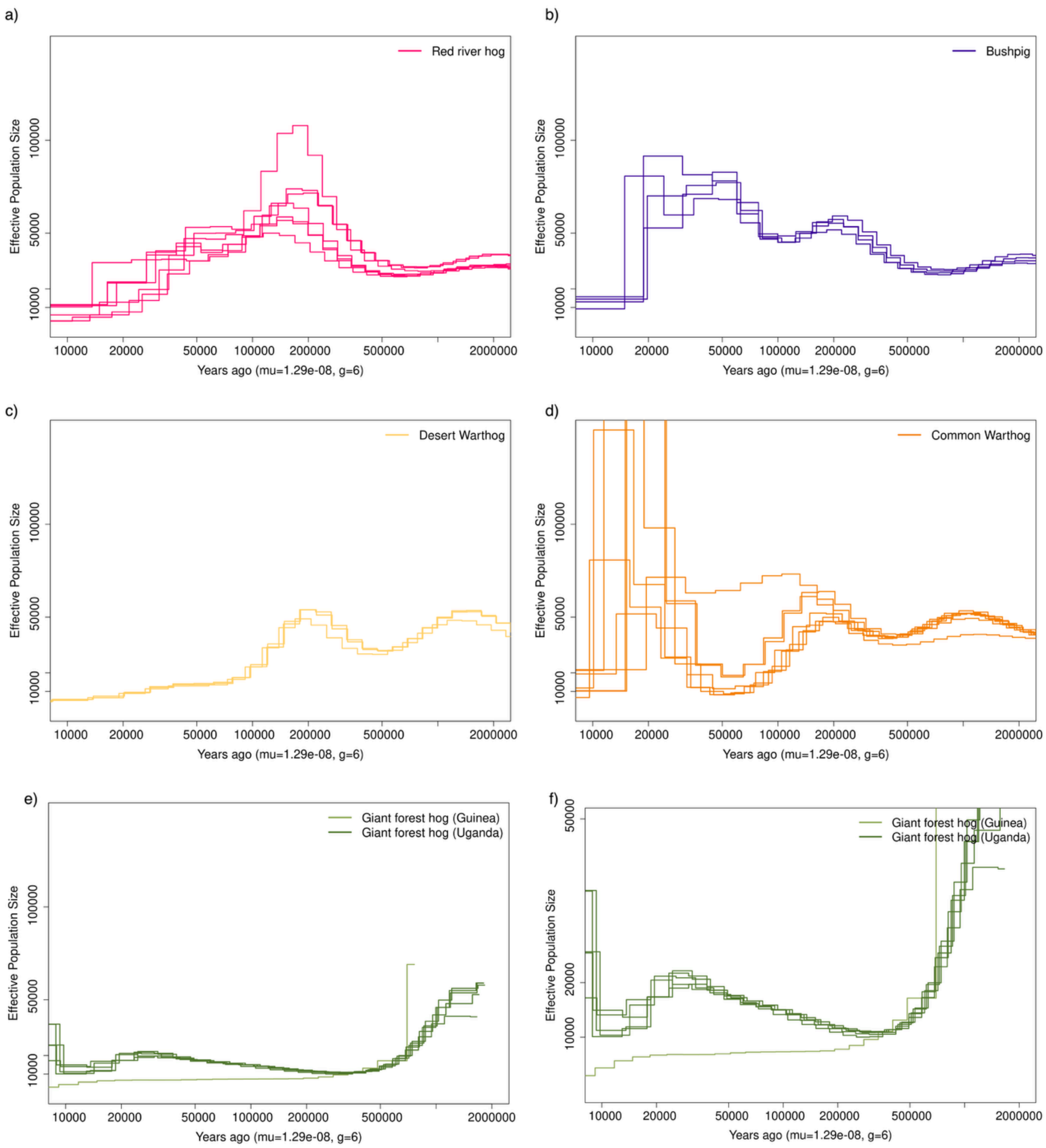


**Figure S3. African suids demographic history inferred with PSMC.**

Demographic history of the a) red river hogs (magenta), b) bushpigs (blue) c) desert warthogs (yellow), d) common warthogs (orange) and e) giant forest hogs (green and light green for sample from Uganda and Guinea respectively) in this study, inferred with PSMC. In panel f) the two populations of GFHs are represented at a larger scale on the y axis. In panel a) and d) the two samples that show different Ne trajectories compared to the rest are both from West Africa (Ghana).

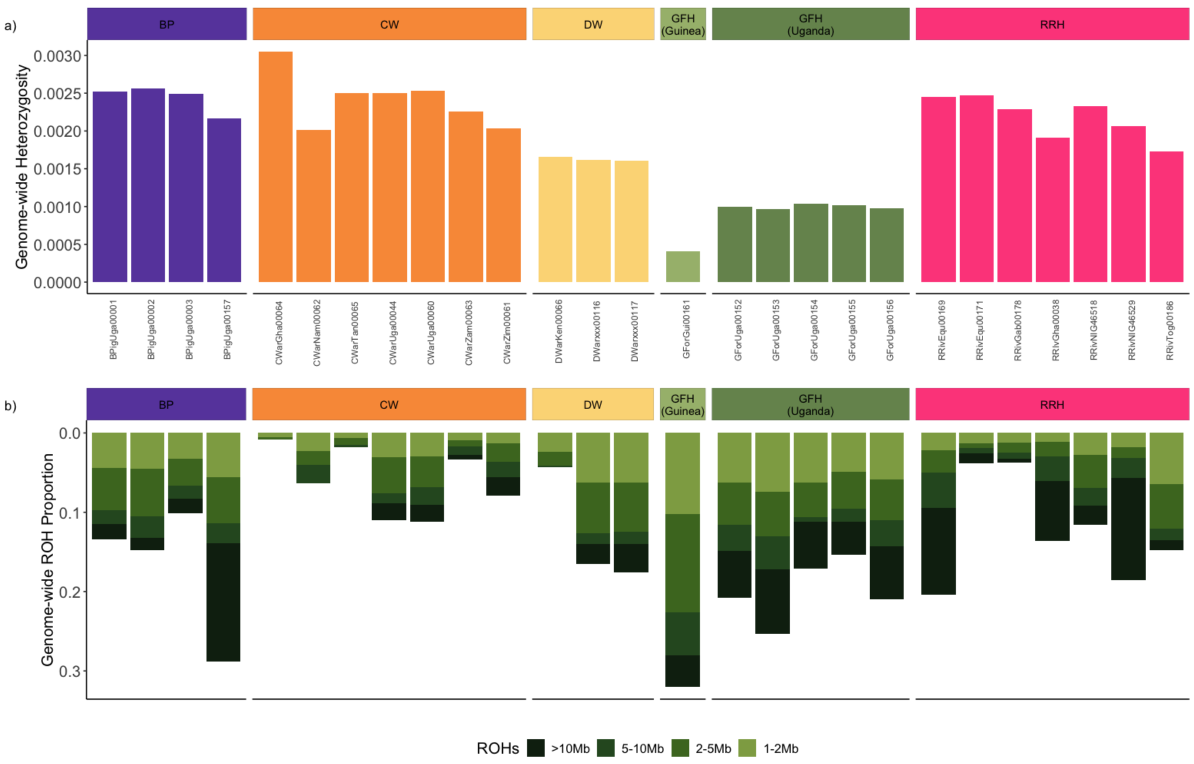


**Figure S4: Genome-wide heterozygosity and ROH proportions.**

a) Genome-wide heterozygosity measured as the proportion of heterozygous sites per bp across each high coverage genome/individual per species. b) Proportions of large (>1MB) genome-wide runs of homozygosity (ROHs) for all high coverage individuals per species. Each individual is represented by a single bar and four sizes of ROHs regions are represented with different shades of green: small sizes are shown in light green while big ROHs are shown in darker green.
